# Using Mapping-Profiles to Refine Strain-Level Metagenomic Classification

**DOI:** 10.64898/2026.05.18.725856

**Authors:** Josipa Lipovac, Lune Angevin, Krešimir Križanović

## Abstract

Metagenomic classification at the strain level remains challenging due to high sequence similarity among closely related genomes, which leads to ambiguous read mappings and frequent false-positive strain detections. Reducing such errors improves the reliability of strain-level analyses, which is critical for applications such as pathogen detection. We introduce StrainRefine, a post-mapping refinement method that analyzes read-reference mapping profiles to resolve ambiguous assignments among highly similar genomes. The method represents candidate reference genomes using binary profiles that capture read-support patterns and measures similarity between references based on profile overlap. The method clusters references based on similar mapping profiles, filters weakly supported genomes, and reassigns reads to representative references, reducing redundant reporting of near-identical strains.

StrainRefine substantially reduces false-positive strain detections while preserving recall and improving agreement between predicted and true abundance profiles. On large-scale metagenomic datasets, it achieves a substantially improved precision–recall balance compared to existing mapping-based approaches, with the standalone method obtaining the highest read-level classification accuracy on the most complex evaluated dataset. Unlike many strain-level tools designed for individual species, StrainRefine operates without prior assumptions about sample composition or curated species-specific reference collections, while still achieving comparable performance in single-species settings on species-specific reference databases. These results highlight mapping-profile similarity as an effective signal for improving strain-level metagenomic classification.

## 1 Introduction

Metagenomics enables the study of microbial communities directly from environmental samples without the need for cultivation. However, classification of metagenomic sequences becomes increasingly challenging at lower taxonomic ranks. In particular, strain-level classification remains difficult due to the high sequence similarity among closely related strains, which often leads to ambiguous read assignments and reduced classification accuracy. Reducing false-positive strain detections is important because it improves the reliability of strain-level analyses, which is critical for applications such as pathogen detection.

Many metagenomic classification tools, such as Kraken2 [21] and Centrifuge / Centrifuger [9, 19], rely on k-mer–based strategies that provide high speed and scalability for large datasets. However, these approaches often struggle to distinguish closely related strains because high sequence similarity reduces the discriminatory power of k-mer–based signals [16].

Mapping-based approaches, instead, align reads to reference genomes and leverage longer sequence context, generally providing higher precision for strain-level identification [14]. However, these methods are computationally more demanding, particularly when large reference databases containing thousands of closely related genomes must be searched. Several methods have been proposed for strain-level analysis. Tools such as StrainScan [11] and StrainGE [2] were primarily developed for short-read datasets and rely on k-mer–based or variant-based signatures to distinguish closely related strains. Mapping-based approaches such as PathoScope [5, 7] instead perform probabilistic reassignment of reads among candidate references based on alignment scores. Long-read sequencing technologies, including Oxford Nanopore and PacBio HiFi, provide longer sequence context that can improve strain-level resolution in metagenomic samples [8, 3].

More recent methods aim to enable strain-level classification in complex multi-species metagenomic datasets and to support long-read sequencing data. For example, ORI [18] and PanTax [23] support strain-level analysis across multiple species, while approaches such as MORA [24] and AugPatho [5, 7, 24] combine read mapping with statistical reassignment to resolve ambiguous mappings. Nevertheless, these methods can become computationally demanding when applied to very large reference databases. The MADRe pipeline [12] introduced an assembly-driven framework for strain-level metagenomic classification that combines database reduction with mapping-based read classification. Although MADRe substantially improves strain-level accuracy, challenges remain when reference databases contain large numbers of closely related genomes, which can lead to redundant or false-positive strain detections.

To address this limitation, we propose StrainRefine, a method that analyzes read–reference mapping profiles to identify clusters of highly similar genomes and leverage this structure to improve read classification and reduce false positive identifications. By combining profile-based similarity analysis, threshold-based filtering, and cluster-driven representative selection, the method reduces redundant strain detections and improves classification precision while maintaining high recall.

The method operates on read–reference mapping results and can therefore be applied as a post-processing step in mapping-based strain-level classification pipelines. In this work, we evaluate Strainrefine both within the MADRe framework and as a standalone refinement step applied to read–reference mappings.

## 2 Methods

### 2.1 Overview of the classification framework

We propose StrainRefine, a method for strain-level metagenomic classification that analyzes relationships between candidate reference genomes using read–reference mapping profiles. The method exploits similarities in read-support patterns across reference genomes to identify groups of highly similar references and refine read assignments. By leveraging this structure, StrainRefine reduces redundant detections of closely related genomes and mitigates false-positive strain identifications while preserving the underlying signal of the true strain present in the sample.

An overview of the proposed method is illustrated in Fig. 1.

**Fig. 1.**
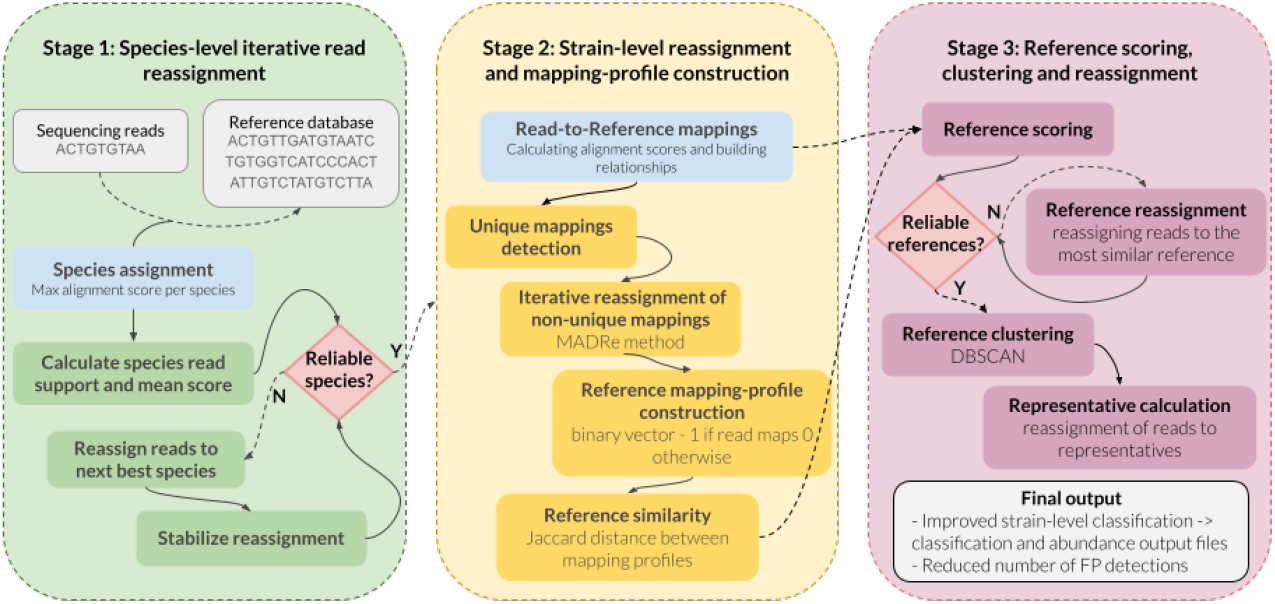
Overview of the StrainRefine workflow. In Stage 1, reads are first assigned to species based on maximum alignment scores. Species-level assignments are iteratively refined by evaluating species read support and mean alignment scores, where reads assigned to weakly supported species are reassigned to better-supported alternatives when available. In Stage 2, reads are classified at the strain level using the MADRe probabilistic reassignment procedure. Read–reference mapping relationships are then represented as binary mapping profiles indicating whether a read maps with the maximum score to a given reference genome. Pairwise similarities between mapping profiles are computed using Jaccard distance. In Stage 3, references with weak read support are identified and their reads are reassigned to the most similar reliable references. Highly similar references are subsequently clustered using DBSCAN, and reads assigned to references within the same cluster are reassigned to representative genomes. The final output consists of refined strain-level classifications with reduced redundant and false-positive strain detections.

StrainRefine operates as a post-mapping refinement step and can be applied to mapping-based classification pipelines that produce read–reference alignments in PAF format.

MADRe [12] framework consists of two main stages: database reduction and read classification. During database reduction, metagenomic assembly is performed and the resulting contigs are mapped to reference genomes, followed by expectation–maximization (EM)-based reassignment of mappings to construct a reduced database of candidate references. This step substantially decreases the size of the reference database while retaining genomes that may be present in the sample, although some false-positive references may remain.

In the read-classification stage, reads are mapped to the reduced database, assigned to species, and then probabilistically reassigned among strain genomes within each species. However, due to mapping ambiguity among closely related genomes, this process can still produce false-positive strain detections. Taking advantage of MADRe’s modular design, we also evaluate multiple read-classification methods, including StrainRefine, MORA, and AugPatho, as drop-in replacements for MADRe’s read-classification stage following database reduction.

Compared to original MADRe’s read-classification step, in StrainRefine, this stage is extended with an additional refinement step that analyzes read–reference mapping profiles to identify relationships among candidate references. These relationships are then used to filter weakly supported references and consolidate signals from highly similar genomes, improving the precision of strain-level classification while maintaining high recall. The individual components of the method are described in the following subsections.

### 2.2 Species-level iterative read reassignment

Species-level assignments are first refined using an iterative reassignment procedure based on read-reference alignment scores. Let *AS*_*r,g*_ denote the normalized alignment score between read *r* and reference genome *g*, computed as

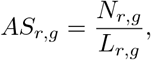

where *N*_*r,g*_ denotes the number of matching bases and *L*_*r,g*_ denotes the mapping length between read *r* and reference genome *g*. Let *G*_*s*_ denote the set of reference genomes belonging to species *s*. For each species *s*, the species score of read *r* is defined as

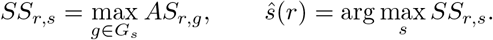

For each species *s*, we compute the number of assigned reads

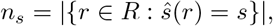

and the mean species score

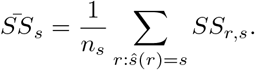

A species is considered unreliable if *n*_*s*_ *<* 5 or if (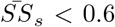 and *n*_*s*_ *<* 30). Reads assigned to unreliable species are reassigned to the alternative species with the highest species score, with ties resolved by selecting the species supported by the largest number of reads. If no reliable alternative species assignment exists, the read remains assigned to the original species to avoid discarding potentially valid low-support signals.

The reassignment procedure is repeated until assignments stabilize, meaning no further changes in read assignments occur, or a maximum of 100 iterations is reached.

This step removes weakly supported species assignments and stabilizes species-level read grouping prior to strain-level classification.

### 2.3 Strain-level probabilistic reassignment

After stabilizing species-level assignments, strain-level classification is performed using the read reassignment procedure previously introduced in MADRe [12]. In StrainRefine, this component is applied without modification to obtain refined strain-level assignments prior to the construction of reference mapping profiles. The following steps correspond to the original procedure. Within each detected species, reads are mapped to candidate reference genomes. Reads that map uniquely to a single genome are directly assigned and remain fixed during reassignment. Reads mapping to multiple candidate genomes are considered ambiguous and are reassigned according to the overall alignment support accumulated by candidate references.

Candidate references are grouped according to shared read mappings, such that references connected through common mapped reads form the same reassignment set. For each reassignment set, total support is computed from both uniquely and ambiguously mapped reads. Ambiguous reads associated with a given set are then reassigned to the reference genome with the strongest overall support within that group.

This procedure consolidates ambiguous mappings toward the most strongly supported references while preserving uniquely supported assignments. The resulting strain-level assignments are subsequently used to construct reference mapping profiles in the following stage.

### 2.4 Reference mapping profiles and within-species distance calculation

After obtaining first-stage strain-level read assignments, binary mapping profiles are constructed for all candidate reference genomes. These profiles represent the set of reads supporting each reference genome and are used to quantify similarity between references belonging to the same species.

For species *s*, let *G*_*s*_ = {*g*_1_, …, *g*_*m*_} denote the candidate reference genomes and *R*_*s*_ = {*r*_1_, …, *r*_*n*_} the reads assigned to that species. Each reference genome *g*_*i*_ is represented by a binary mapping profile 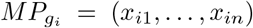, where *x*_*ij*_ = 1 if read *r*_*j*_ maps with maximum score to *g*_*i*_ and 0 otherwise.

Pairwise Jaccard distances [10] are then computed between mapping profiles of references within the same species in order to quantify similarity between their read-support patterns.

A similar comparison of mapping patterns across highly similar genomes was briefly explored in the MADRe study for evaluation purposes. In contrast, in this work mapping-profile distances are used as a core component of the proposed method.

### 2.5 Reference scoring and threshold-based reassignment

After constructing mapping profiles, candidate reference genomes are evaluated based on the strength of their read support.

For each reference genome *g*, a normalized mapping score in the range [0, 1] is computed using the alignment score defined in Section 2.2, where higher values indicate stronger support for the corresponding reference genome. The score reflects the relative confidence of assigning a read to a given reference and is identical to the scoring scheme used in MADRe [12]. Based on these scores, we count the total number of mapped reads and the number of reads with alignment scores *>* 0.6.

Let *h*_*g*_ denote the number of reads mapped to reference *g* with alignment scores *>* 0.6. A reference is considered unreliable if *h*_*g*_ *<* 5, or if (5≤ *h*_*g*_ *<* 10 and *GS*_*g*_ *<* 0.8), where *GS*_*g*_ is the mean mapping score of reads mapped to *g*.

Reads assigned to unreliable references are reassigned to the most similar reliable reference according to mapping-profile distances. If no sufficiently similar reliable reference exists, the reads remain assigned to the original reference to avoid incorrect reassignment when reads originate from low-abundance or unrepresented genomes.

### 2.6 Cluster-based representative selection and read reassignment

Reference genomes belonging to the same species are clustered based on pairwise Jaccard distances between their mapping profiles. Clustering is performed independently for each species using the DBSCAN [4] algorithm applied to the mapping-profile distance matrix.

The neighborhood radius parameter *eps* was determined empirically through grid search on simulated datasets used during method development, with the minimum number of samples set to 1. The final value *eps* = 0.8 was selected based on analysis of average nucleotide identity (ANI) between genomes grouped within clusters and between genomes belonging to different clusters, ensuring high similarity within clusters while maintaining clear separation between them. For each cluster, a representative reference genome is selected as the genome with the largest number of reads mapped to it prior to the final reassignment step. Reads assigned to references within the same cluster are then reassigned to the corresponding representative genome, thereby merging detections of highly similar references while preserving the overall signal of the underlying strain. References that belong to single-genome clusters remain unchanged and retain their original read assignments.

### 2.7 Parameter selection and sensitivity analysis

Default parameter values were selected empirically during method development using sensitivity analyses on representative datasets. Key parameters controlling clustering behavior and reference/species reliability filtering were evaluated across a range of values, including the DBSCAN clustering threshold (*ϵ*), alignment score thresholds, minimum support requirements, and species-level reassignment criteria. Default values were chosen to provide a balance between reducing false-positive detections and maintaining read-level classification performance.

Sensitivity analyses for all evaluated parameters and selected default values are provided in Supplementary File (Supplementary Fig. S1).

## 3 Results

### 3.1 Experimental setup

The performance of StrainRefine was evaluated in two experimental scenarios designed to assess both large-scale metagenomic classification and strain-level resolution in single-species datasets. A summary of the experimental setups is provided in Table 1. All sequencing reads used in the experiments, including simulated datasets and the Zymo mock community, correspond to Oxford Nanopore Technologies (ONT) R10 reads.

**Table 1.** Summary of experimental scenarios used for evaluating StrainRefine.

| Scenario | Dataset | # | Database | Compared tools |
| --- | --- | --- | --- | --- |
| Large-scale classification | 66s (PanTax) | 66 | NCBI RefSeq | MADRe+SR, SR, MADRe, MADRe_RC, MORA, AugPatho |
| Large-scale classification | Zymo D6331 | 31 (5 *E.coli*) | MADRe reduced | SR, MADRe_RC, MORA, AugPatho |
| Large-scale classification | 1000s (Pantax) | 1063 | NCBI RefSeq | MADRe+SR, SR, MADRe, MADRe_RC, MORA, AugPatho |
| Single-species resolution | *P.aeruginosa* (synthetic) | 3 | MADRe-reduced | SR, MADRe_RC, StrainGE, StrainScan, ORI |
| Single-species resolution | *P.aeruginosa* (synthetic) | 5 | MADRe-reduced | SR, MADRe_RC, StrainGE, StrainScan, ORI |
| Single-species resolution | *P.aeruginosa* (synthetic) | 10 | MADRe-reduced | SR, MADRe_RC, StrainGE, StrainScan, ORI |

In the first scenario, StrainRefine was evaluated both as an integrated component of the MADRe pipeline (MADRe+SR) and as a standalone post-processing method (SR) applied to read–reference mapping results. The evaluation also included the original MADRe pipeline and its standalone read-classification step (MADRe_RC), together with other mapping-based strain-level classification tools capable of handling large reference databases, including MORA and Aug-Patho. Unless otherwise stated, all methods were evaluated on metagenomic datasets using a comprehensive NCBI reference database corresponding to the same NCBI RefSeq release used in the original MADRe study (December 2024) [15].

Experiments in this scenario were performed using the 66s dataset containing 60 strains represented by 66 reference sequences [22] and the high-complexity dataset from the PanTax benchmark [22], comprising 1,063 reference sequences spanning over 300 species and inspired by the CAMI challenge design [6]. Simulated ONT R10 reads for the high-complexity dataset were generated in the original MADRe study [12] using the published reference genomes and expected abundances with the Badread tool [20].

We additionally evaluated the Zymo D6331 mock community dataset [13], which contains real ONT sequencing reads from 18 bacterial strains (non-bacterial organisms were excluded) represented by 31 reference sequences, as some strains consist of multiple genomic elements (e.g., chromosomes or plasmids). This dataset includes five strains of *Escherichia coli*, providing a challenging setting with highly similar strains and allowing us to specifically evaluate the behavior of reassignment methods under substantial mapping ambiguity.

For this dataset, the evaluation focused specifically on the reassignment stage. To reduce the impact of large-scale database search, reads were first processed through the MADRe database reduction step, and candidate references from the reduced database were subsequently used as database for SR, MADRe_RC, MORA, and AugPatho. Although the database was reduced, it still contained 316 *E. coli* reference genomes, preserving substantial strain-level similarity and ambiguity while reducing the overall search space.

Throughout the evaluation, detections correspond to unique reference accessions present in the reference FASTA files. In some cases, multiple accessions represent genomes originating from the same biological strain; however, evaluating predictions at the accession level provides a direct and reproducible comparison with the reference database used for classification.

In the second scenario, StrainRefine (SR) was evaluated as a standalone method applied to read–reference mapping results. As in the first scenario, reference database was first reduced using the MADRe database reduction procedure. This reduction was necessary because several strain-level identification tools operate on relatively small reference collections, often restricted to a single species, and cannot efficiently process very large reference databases.

These experiments focused on datasets containing strains from a single species to evaluate the ability to resolve highly similar reference genomes. Synthetic datasets containing 3, 5, and 10 strains of *Pseudomonas aeruginosa* were generated using the Badread simulator with equal abundances (20× coverage). *P. aeruginosa* was selected because the reference database contains a large number of closely related strains, providing a suitable test case for evaluating false-positive reduction among highly similar genomes. Genomes were selected to cover different ranges of pairwise similarity according to average nucleotide identity (ANI). The list of genomes and their pairwise ANI values are provided in the Supplementary Material.

For comparison in the single-species resolution experiments, we included several strain-level identification tools designed for smaller reference collections. We evaluated the original MADRe read-classification procedure (MADRe_RC), ORI, and two methods originally developed for short-read data, StrainScan and StrainGE, which can also be applied to long-read sequencing datasets. Although we initially planned to include PanTax in this comparison, repeated attempts to construct the required reference database were unsuccessful, and PanTax was therefore excluded from the final evaluation.

### 3.2 Large-scale classification results

Results on the 66s dataset highlight clear differences in the precision–recall trade-off between the evaluated tools. As shown in the precision–recall plot (Fig. 2, top), MADRe+SR achieves substantially higher precision than the other methods, including the original MADRe, while maintaining high recall. Strain-level results (Fig. 2, tables) show that MADRe+SR misses only one strain detected by MADRe. After clustering highly similar genomes, following the evaluation strategy used in the original MADRe study, the missing strain is merged with a nearly identical reference, largely eliminating the difference between the methods. Overall, MADRe+SR reports a number of strains close to the ground-truth count, while some low-abundance strains remain undetected.

**Fig. 2.**
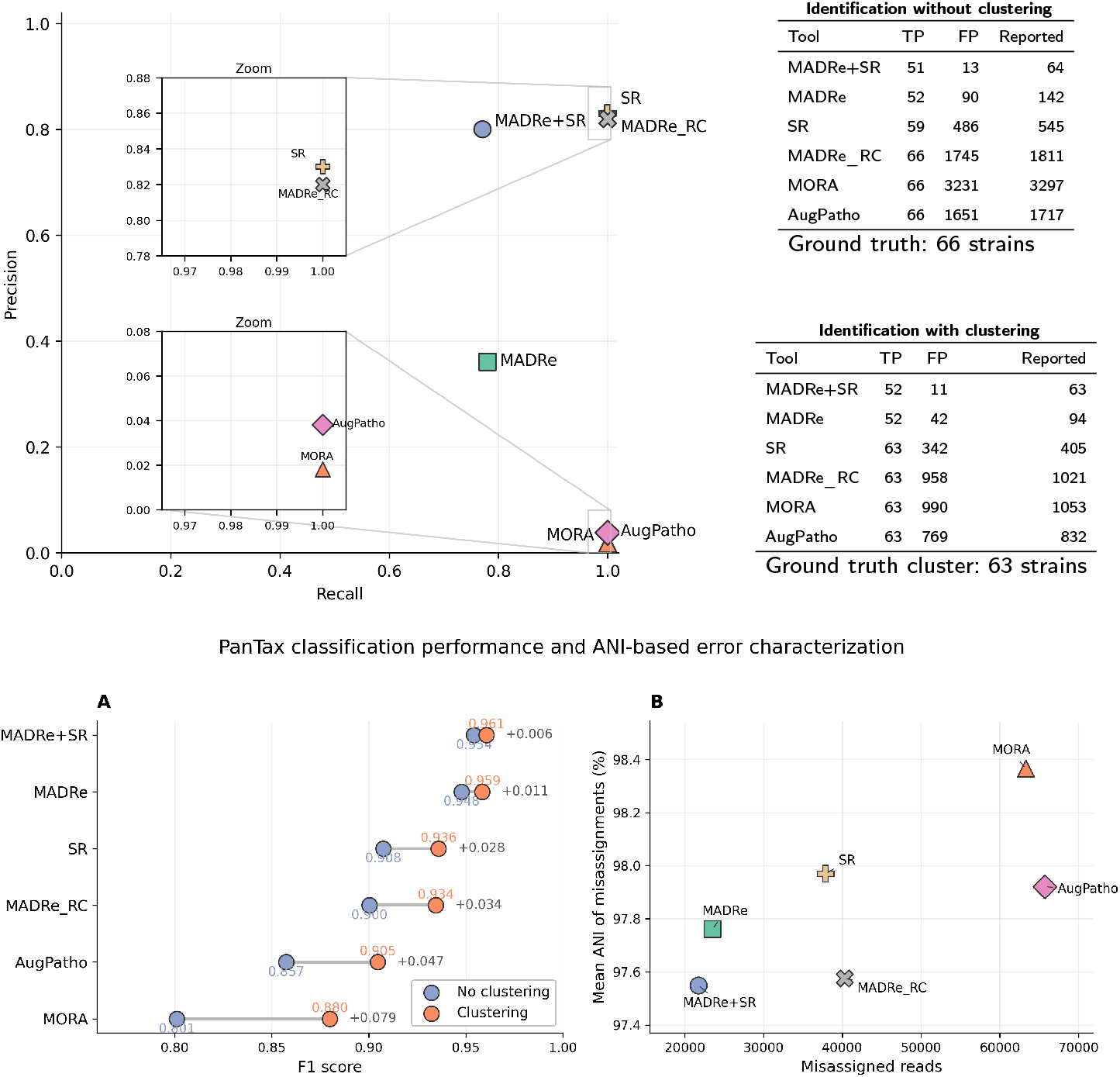
Evaluation of strain identification and read assignment accuracy on the 66s dataset. **Top:** Precision-recall comparison of reported strain calls across methods (inset: high-recall region). Tables summarize true positives (TP), false positives (FP), and reported accessions before and after clustering highly similar genomes (66 strains*→* 63 clusters). **Bottom:** Read-level error analysis: (A) F1 scores before and after clustering;(B) relationship between the number of misassigned reads and their mean ANI.

The standalone SR approach and MADRe_RC achieve similar overall precision– recall performance; however, their strain-level identification profiles differ substantially. Without clustering, SR reports considerably fewer strain identifications than MADRe_RC (545 vs. 1811), resulting in substantially fewer false-positive detections. Many of the removed detections correspond to references supported by a very small numbers of reads. Consequently, the improvement in overall precision and F1 score remains moderate despite the large reduction in reported strains. Although SR initially appears to miss several strains before clustering, these strains belong to the same clusters as the predicted references and are therefore effectively recovered after clustering highly similar genomes. After clustering, SR achieves precision and recall values close to MADRe_RC while still reporting substantially fewer false-positive identifications.

The read-level analysis (Fig. 2, bottom) further illustrates the benefit of this reduction in reported strains. MADRe+SR achieves the highest F1 score, with substantially fewer false-positive read assignments than competing tools. SR also reduces the number of misassigned reads compared to MADRe_RC, and the remaining misassignments occur between genomes with higher mean ANI values, indicating that the residual classification errors are more frequently restricted to highly similar strains. In contrast, MORA and AugPatho produce substantially larger numbers of misassigned reads despite many of these errors also occurring between closely related genomes.

Importantly, the performance of MADRe+SR remains nearly unchanged before and after clustering highly similar genomes, indicating that the method correctly identifies the underlying biological strains even in the presence of many near-identical references.

To evaluate scalability to higher-complexity communities, we tested all methods on a dataset comprising 1,063 reference genomes (Fig. 3). Since MADRe_RC corresponds to the read-classification and reassignment component of the MADRe pipeline, and SR can be applied as a direct replacement for this step, a comparison between these two methods directly quantifies the effect of StrainRefine refinement. SR reduces false-positive organism identifications from 18,734 to 5,238 (approximately 72%) while maintaining a nearly identical number of true positives (1,011 vs. 1,049), and achieves the highest read-level F1 score among all evaluated methods (0.912).

**Fig. 3.**
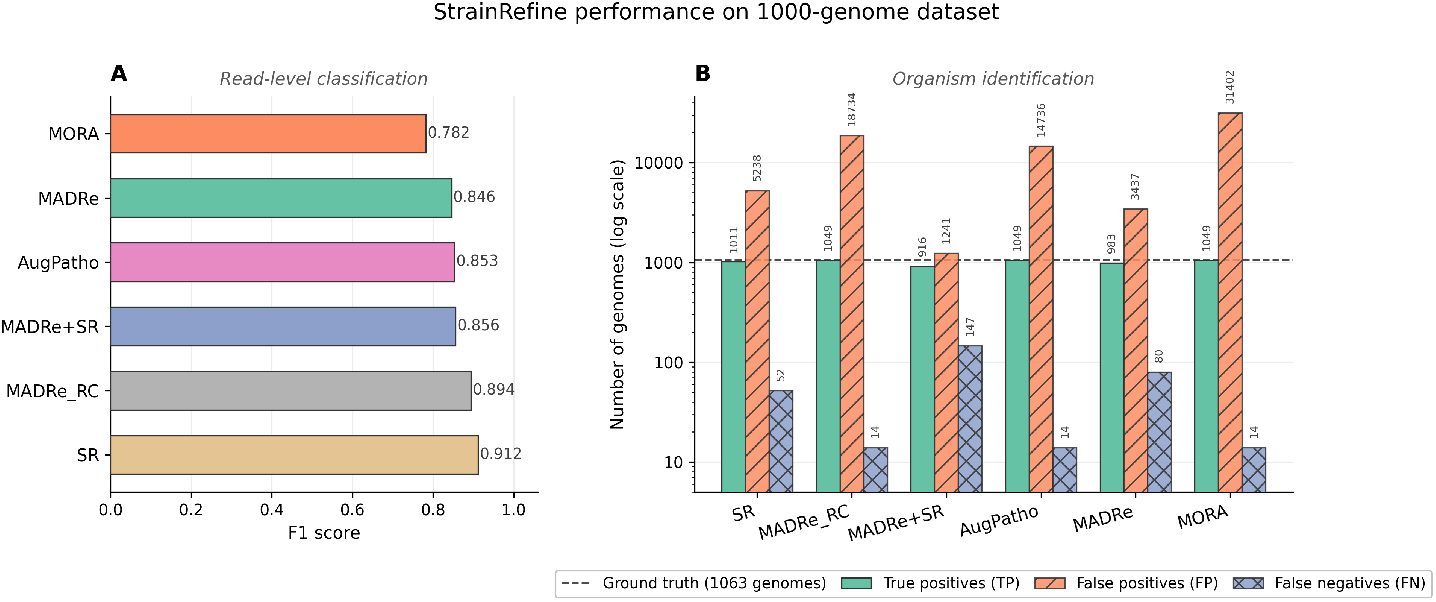
Classification performance on the 1000-genome dataset. **(A)** Read-level classification accuracy (F1 score) for all evaluated methods. **(B)** Organism identification performance showing true positives (TP), false positives (FP), and false negatives (FN); y-axis shown in log scale. The dashed line indicates the ground truth of 1,063 reference genomes.

When StrainRefine is fully integrated into the MADRe pipeline (MADRe+SR), the number of false positives decreases further to 1,241, demonstrating that the combination of assembly-driven database reduction and refinement provides the strongest overall precision–recall balance. In contrast, methods without an explicit refinement stage, such as MORA and MADRe_RC, exhibit substantially higher false-positive counts as dataset complexity increases.

The apparent discrepancy between read-level F1 scores and organism-level false-positive counts is explained by the nature of false detections: most falsely reported organisms are supported by only a very small number of reads, contributing minimally to read-level metrics while substantially inflating organism-level identification counts.

To assess the behavior of reassignment methods on the Zymo dataset after database reduction, we computed the Bray–Curtis distance between predicted and ground-truth abundance profiles (**Fig. 4A**) and evaluated the number of false positive identifications (**Fig. 4B**). SR achieved the lowest Bray–Curtis distance for the complete community profile and for the non-*E. coli* subset, indicating the strongest overall agreement with the true abundance distribution. In the *E*.*coli* -only subset, which contains multiple highly similar strains and represents the most challenging part of the dataset, AugPatho achieved the lowest Bray– Curtis distance, while SR remained competitive.

**Fig. 4.**
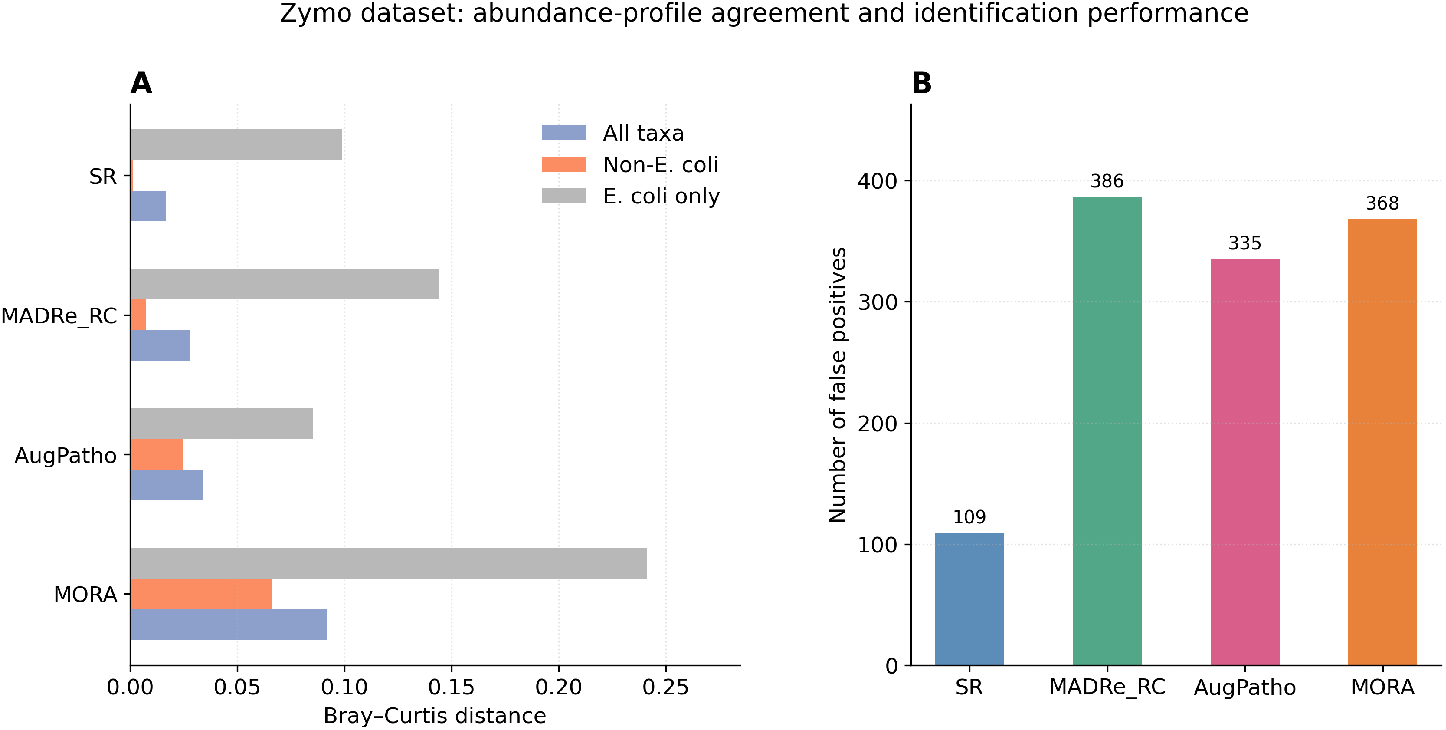
Strain-level classification results on the Zymo dataset. **(A)** Bray–Curtis distance between predicted and true abundance profiles (all taxa, non-*E. coli*, and *E. coli* strains; lower is better). **(B)** Identification performance in number of FPs.

SR also substantially reduced the number of false positive strain identifications compared to the other evaluated reassignment methods. Specifically, SR reported 109 false positives, compared to 386 for MADRe_RC, 335 for Aug-Patho, and 368 for MORA, corresponding to more than a threefold reduction relative to MADRe_RC.

### 3.3 Single-species resolution results

In the single-species resolution experiments, we assessed the performance of SR on reduced single-species reference databases to determine whether it can reduce false positive (FP) identifications while remaining competitive with tools specifically designed for strain-level analysis within individual species. The results across datasets of increasing complexity are summarized in Figure 5. Across all evaluated datasets, SR consistently detected all expected strains while maintaining a low number of FP predictions. In particular, the reduction in FP counts compared to the original MADRe read-classification procedure is substantial across all tested experimental setups. Detailed quantitative results, including abundance estimates for individual strains, are provided in Supplementary Table 2.

**Fig. 5.**
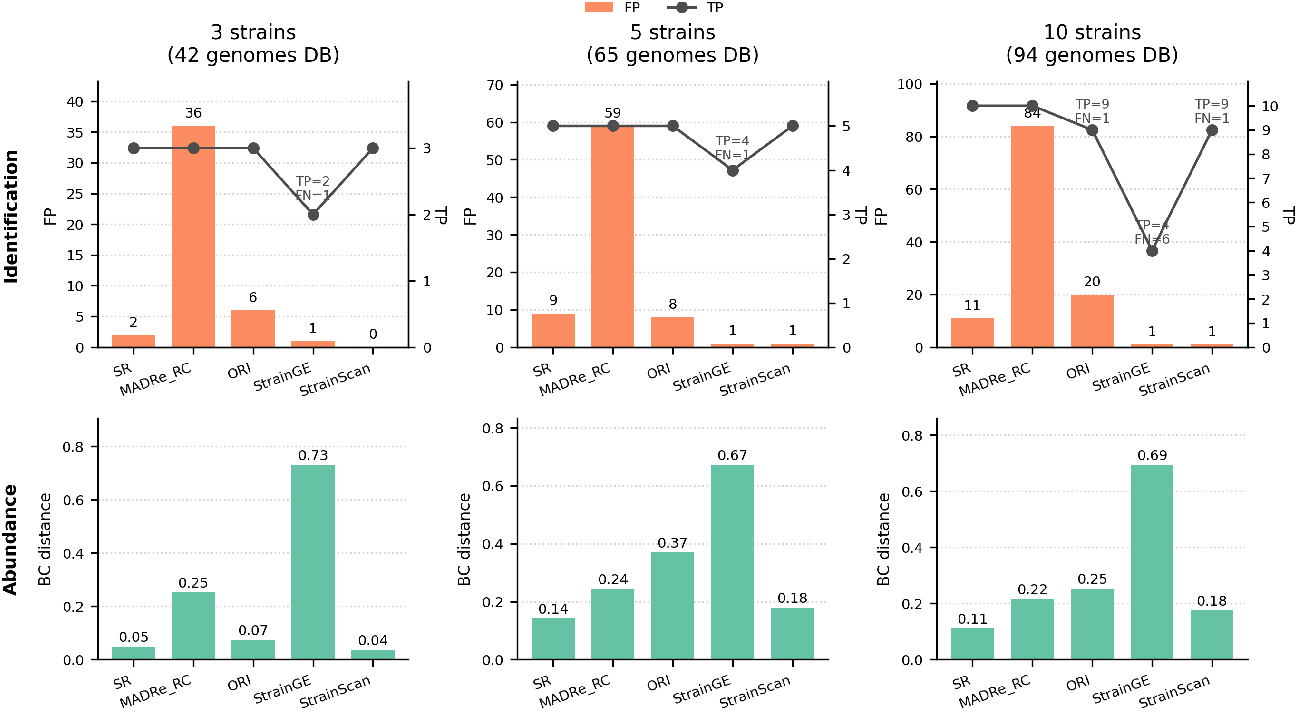
Strain-level identification and abundance estimation on single-species datasets of increasing complexity. Top: False positives (FP; bars) and true positives (TP; line); false negatives (FN) annotated. Bottom: Bray–Curtis (BC) distance between predicted and ground-truth abundance profiles (lower is better). Columns correspond to datasets with 3, 5, and 10 strains and reduced reference databases of 42, 65, and 94 genomes.

Among the evaluated methods, StrainScan achieves the lowest FP counts while correctly detecting all expected strains. However, the Bray–Curtis distances indicate that SR provides more accurate abundance estimates, showing closer agreement with the ground-truth abundance profiles across all datasets. ORI also produces relatively low FP counts, but begins to miss expected strains as the dataset complexity increases, particularly in the datasets containing larger numbers of highly similar genomes.

It is important to note that the evaluated single-species tools internally perform various forms of clustering or grouping of highly similar reference genomes as part of their classification procedures. Similarly, SR groups candidate references based on mapping-profile similarity. Therefore, we did not apply any additional clustering during evaluation, allowing the results to reflect how each method selects representative strains among highly similar references.

### 3.4 Discussion

Strain-level metagenomic classification remains challenging because closely related reference genomes often produce highly ambiguous read mappings. Reads originating from the same strain may map equally well to several references, causing mapping-based classification pipelines to report multiple highly similar genomes. This leads to redundant detections and large numbers of false-positive strain calls. In this work, we introduce StrainRefine, a method that analyzes read–reference mapping profiles to identify clusters of highly similar genomes and refine read classification to reduce false-positive strain identifications.

When integrated into the MADRe pipeline as a replacement for the MADRe_RC step, StrainRefine substantially reduced false-positive strain detections while preserving competitive classification performance. Since read mapping represents the most computationally demanding step in mapping-based metagenomic classification, as observed in the original MADRe study, combining StrainRefine with MADRe enables refinement to operate on a reduced reference database, thereby limiting additional computational cost. At the same time, StrainRefine also achieves strong classification performance as a standalone method and substantially reduces false-positive detections compared to MADRe_RC.

Sensitivity analyses suggest that StrainRefine is generally robust to moderate parameter variation. Most evaluated thresholds produced relatively stable performance across a broad range of values, indicating that the method is not highly dependent on precise parameter tuning. The largest sensitivity was observed for the clustering threshold (*ϵ*), where extreme values can either separate highly similar references into distinct groups or merge biologically distinct clusters. Other parameters primarily influenced the trade-off between false-positive reduction and read-level sensitivity. Detailed parameter analyses are provided in Supplementary File.

Results on the 66s and high-complexity datasets demonstrated a consistent pattern across increasing levels of community complexity. On the 66s dataset, MADRe+SR achieved an improved precision–recall balance compared to the original MADRe pipeline and other evaluated methods, while also improving read-level classification accuracy. Similar trends were observed on the high-complexity dataset, where SR achieved the highest read-level F1 score when applied as a standalone method. However, this configuration requires mapping reads against the full reference database, which constitutes the most computationally expensive stage of mapping-based pipelines. When StrainRefine is instead applied after the MADRe database reduction step, computational requirements are substantially reduced while maintaining competitive classification performance. The choice between the two configurations therefore depends on the available computational resources and the relative priority of read-level accuracy versus organism-level precision.

Results on the Zymo mock community dataset further highlight the behavior of different reassignment strategies in the presence of highly similar strains and real sequencing data. Since the dataset contains multiple *E. coli* strains and the reduced database still retained 316 *E. coli* reference genomes, substantial ambiguity remained despite database reduction. Under these conditions, SR achieved the lowest Bray–Curtis distance for the complete community profile and the non-*E. coli* subset, indicating stronger overall agreement with the true abundance distribution. Although AugPatho achieved slightly better agreement within the *E. coli* -only subset, SR substantially reduced false-positive strain detections compared to all evaluated methods, achieving more than a threefold reduction relative to MADRe_RC.

In the single-species resolution experiments, SR consistently reduced false positives compared to the original MADRe read-classification procedure while maintaining detection of all expected strains. The Bray–Curtis distances indicate improved agreement with the ground-truth abundance profiles, suggesting more accurate abundance estimation even in datasets containing many highly similar genomes. While some specialized strain-level tools can achieve very low false-positive rates in such settings, this may come at the cost of reduced recall as dataset complexity increases. In contrast, SR maintained detection of all expected strains while substantially reducing false positives relative to the original MADRe classification step.

Conceptually, SR also groups highly similar reference genomes, but constructs these groups differently from the compared single-species tools. Methods such as StrainScan and StrainGE cluster genomes based on sequence similarity within predefined reference collections, while ORI collapses nearly indistinguishable strains using genomic-distance criteria. In contrast, SR derives groups directly from read–reference mapping profiles observed in the sample, allowing clustering to reflect the signal present in the data rather than reference similarity or assembly artifacts. When distinguishing regions are weakly covered, clusters may become larger but the idea is that they still reflect the signal supported by the data.

Another potentially useful signal for reducing false-positive identifications is horizontal genome coverage, which has been successfully applied in methods such as SLIMM [1] and KMCP [17]. Since StrainRefine operates on read–reference mappings in PAF format, genome coverage profiles can be readily computed from mapped read coordinates. However, incorporating fixed horizontal coverage thresholds into strain-level refinement is not straightforward in the context of highly similar genomes. Reads originating from closely related strains frequently accumulate on conserved genomic regions shared across multiple references, while strain-specific regions may receive only sparse or uneven support. As a result, true-positive strains, particularly those with low abundance, may exhibit limited horizontal coverage despite being genuinely present in the sample. In this setting, low genome breadth does not necessarily correspond to false detection. For this reason, StrainRefine currently relies primarily on relationships between read-support patterns rather than explicit coverage thresholds, although genome coverage information could represent a useful complementary signal in future extensions of the framework.

The current formulation of StrainRefine is primarily designed for long-read sequencing data, as the proposed mapping-profile analysis benefits from the longer sequence context provided by long reads. Although long-read technologies typically exhibit higher sequencing error rates, the extended genomic context captured by individual reads still provides more informative and discriminative read–reference mapping relationships across highly similar genomes. This richer contextual signal improves the ability to identify consistent mapping-support patterns between related references.

Future work will evaluate the method on more complex communities and explore detection of novel sequences absent from the reference database. Overall, these results suggest that analyzing similarities between reference mapping profiles provides a useful signal for resolving ambiguous assignments among closely related genomes and improving strain-level metagenomic classification.

## Acknowledgments

This work was supported by the Croatian Science Foundation under grants IP-2018-01-5886 (SIGMA) and DATACROSS (2024–2026, PK.1.1.10.0007).

The implementation of StrainRefine is available at: https://github.com/lbcb-sci/StrainRefine.

## Disclosure of Interests

The authors have no competing interests to declare.

## Notes

### Competing Interest Statement

The authors have declared no competing interest.

## References

1. Dadi, T.H., Renard, B.Y., Wieler, L.H., Semmler, T., Reinert, K.: Slimm: species level identification of microorganisms from metagenomes. PeerJ 5, e3138 (2017)

2. van Dijk, L.R., Walker, B.J., Straub, T.J., Worby, C.J., Grote, A., Schreiber IV, H.L., Anyansi, C., Pickering, A.J., Hultgren, S.J., Manson, A.L., et al.: Strainge: a toolkit to track and characterize low-abundance strains in complex microbial communities. Genome biology 23(1), 74 (2022)

3. Dilthey, A.T., Jain, C., Koren, S., Phillippy, A.M.: Strain-level metagenomic assignment and compositional estimation for long reads with metamaps. Nature communications 10(1), 3066 (2019)

4. Ester, M., Kriegel, H.P., Sander, J., Xu, X., et al.: A density-based algorithm for discovering clusters in large spatial databases with noise. In: kdd. vol. 96, pp. 226–231 (1996)

5. Francis, O.E., Bendall, M., Manimaran, S., Hong, C., Clement, N.L., Castro-Nallar, E., Snell, Q., Schaalje, G.B., Clement, M.J., Crandall, K.A., et al.: Pathoscope: species identification and strain attribution with unassembled sequencing data. Genome research 23(10), 1721–1729 (2013)

6. Fritz, A., Hofmann, P., Majda, S., Dahms, E., Dröge, J., Fiedler, J., Lesker, T.R., Belmann, P., DeMaere, M.Z., Darling, A.E., et al.: Camisim: simulating metagenomes and microbial communities. Microbiome 7(1), 17 (2019)

7. Hong, C., Manimaran, S., Shen, Y., Perez-Rogers, J.F., Byrd, A.L., Castro-Nallar, E., Crandall, K.A., Johnson, W.E.: Pathoscope 2.0: a complete computational framework for strain identification in environmental or clinical sequencing samples. Microbiome 2, 1–15 (2014)

8. Kim, C., Pongpanich, M., Porntaveetus, T.: Unraveling metagenomics through long-read sequencing: a comprehensive review. Journal of translational medicine 22(1), 111 (2024)

9. Kim, D., Song, L., Breitwieser, F.P., Salzberg, S.L.: Centrifuge: rapid and sensitive classification of metagenomic sequences. Genome research 26(12), 1721–1729 (2016)

10. Levandowsky, M., Winter, D.: Distance between sets. Nature 234(5323), 34–35 (1971)

11. Liao, H., Ji, Y., Sun, Y.: High-resolution strain-level microbiome composition analysis from short reads. Microbiome 11(1), 183 (2023)

12. Lipovac, J., Šikić, M., Vicedomini, R., Križanović, K.: Madre: Strain-level metagenomic classification through assembly-driven database reduction. GigaScience p. giag030 (03 2026)

13. Liu, L., Yang, Y., Deng, Y., Zhang, T.: Nanopore long-read-only metagenomics enables complete and high-quality genome reconstruction from mock and complex metagenomes. Microbiome 10(1), 209 (2022)

14. Marić, J., Križanović, K., Riondet, S., Nagarajan, N., Šikić, M.: Comparative analysis of metagenomic classifiers for long-read sequencing datasets. BMC bioinformatics 25(1), 15 (2024)

15. O’Leary, N.A., Wright, M.W., Brister, J.R., Ciufo, S., Haddad, D., McVeigh, R., Rajput, B., Robbertse, B., Smith-White, B., Ako-Adjei, D., et al.: Reference sequence (refseq) database at ncbi: current status, taxonomic expansion, and functional annotation. Nucleic acids research 44(D1), D733–D745 (2016)

16. Schaeffer, L., Pimentel, H., Bray, N., Melsted, P., Pachter, L.: Pseudoalignment for metagenomic read assignment. Bioinformatics 33(14), 2082–2088 (2017)

17. Shen, W., Xiang, H., Huang, T., Tang, H., Peng, M., Cai, D., Hu, P., Ren, H.: Kmcp: accurate metagenomic profiling of both prokaryotic and viral populations by pseudo-mapping. Bioinformatics 39(1), btac845 (2023)

18. Siekaniec, G., Roux, E., Lemane, T., Guédon, E., Nicolas, J.: Identification of isolated or mixed strains from long reads: a challenge met on streptococcus thermophilus using a minion sequencer. Microbial genomics 7(11), 000654 (2021)

19. Song, L., Langmead, B.: Centrifuger: lossless compression of microbial genomes for efficient and accurate metagenomic sequence classification. Genome biology 25(1), 106 (2024)

20. Wick, R.R.: Badread: simulation of error-prone long reads. Journal of Open Source Software 4(36), 1316 (2019)

21. Wood, D.E., Lu, J., Langmead, B.: Improved metagenomic analysis with kraken 2. Genome biology 20, 1–13 (2019)

22. Zhang, W.: Benchmarking datasets used in the manuscript “strain-level metagenomic profiling using pangenome graphs with pantax” (2025). 10.5281/zenodo.16885808, https://zenodo.org/records/16885808, version v3; accessed 2025-10-19

23. Zhang, W., Liu, Y., Li, G., Xu, J., Chen, E., Schönhuth, A., Luo, X.: Strain-level metagenomic profiling using pangenome graphs with pantax. Genome Research 36(2), 405–420 (2026)

24. Zheng, A., Shaw, J., Yu, Y.W.: Mora: abundance aware metagenomic read re-assignment for disentangling similar strains. BMC bioinformatics 25(1), 161 (2024)

