## Supplementary File for "Using Mapping-Profiles to Refine Strain-Level Metagenomic Classification"

### Sensitivity analysis

To characterize the behavior of StrainRefine under different parameter configurations, we performed sensitivity analyses by varying one parameter at a time while keeping all remaining parameters fixed at their default values. Performance was evaluated using strain-level identification metrics, including F1 score, precision (strain F1), recall (strain recall), and the number of false-positive strain detections (FP strains), together with read-level classification performance measured using read F1 score (read F1).

Sensitivity analyses were performed under three complementary evaluation settings designed to capture different sources of ambiguity encountered in strain-level metagenomic classification:

- **High-complexity community setting:** the 1063s dataset evaluated using the MADRe reduced database. This setting was selected to evaluate parameter behavior in highly complex metagenomic communities containing many species and reference genomes while operating under the reduced-database conditions used in the MADRe+SR pipeline.
- **Large-reference-database setting:** the 66s dataset evaluated against the complete NCBI reference database. This setting was selected to evaluate parameter behavior in the presence of large-scale database ambiguity arising from highly redundant reference collections.
- **Single-species resolution setting:** the dataset containing ten strains from a single species evaluated using the MADRe reduced database. This setting was selected to evaluate parameter behavior in local strain-resolution scenarios dominated by highly similar genomes.

The following default parameter values were used throughout all experiments unless otherwise stated:

- DBSCAN clustering threshold $\epsilon$ : 0.8 - controls similarity required for grouping reference mapping profiles into the same cluster

- Alignment score threshold: 0.6 - defines the minimum normalized alignment score required for a mapping to be considered sufficiently supported

- Minimum reads for reliable references: 5 - minimum number of sufficiently supported reads required for a reference to be considered reliable

- Intermediate support threshold: 10 - defines the read-count threshold below which references are treated as moderately supported rather than strongly supported during reassignment

- Minimum fraction for intermediate-support references: 0.8 - for moderately supported references, defines the minimum fraction of sufficiently supported reads required for retaining the reference during reassignment

- Species minimum read count: 5 - minimum number of assigned reads required for a species to avoid being considered unreliable

- Species minimum mean score: 0.6 - minimum average normalized alignment score required for low-read-count species assignments to avoid being considered unreliable

- Species low-count cap: 30 - upper read-count limit below which low-scoring species assignments may still be considered unreliable

The evaluated parameters include clustering criteria, reference reliability thresholds, and species-level filtering thresholds that influence the reassignment and clustering behavior of StrainRefine.

Figures S1–S3 present sensitivity analysis results across the evaluated experimental settings. In each analysis, one parameter was varied at a time while all remaining parameters were fixed at their default values. Vertical dashed lines indicate the default parameter values used throughout the study.

#### DBSCAN clustering threshold ($\epsilon$)

The DBSCAN clustering threshold produced the strongest overall effect on performance across all evaluated settings. Increasing $\epsilon$ generally reduced the number of false-positive strain identifications by allowing reference genomes with less similar mapping profiles to be grouped into the same cluster. This behavior was particularly evident in the high-complexity and large-reference-database settings, where increasing $\epsilon$ progressively reduced the number of reported false-positive strain identifications while improving strain identification precision.

In the large-reference-database setting, performance continued improving up to $\epsilon = 0.9$, suggesting that more permissive clustering may be beneficial when operating on highly redundant reference collections containing many closely related genomes. This likely reflects the increased degree of mapping ambiguity present in large databases, where reads originating from a single strain may support a larger number of highly similar references.

However, extreme values ($\epsilon = 1.0$) substantially reduced strain identification recall and read-level classification performance. In the context of binary mapping-profile clustering using Jaccard distance, $\epsilon = 1.0$ effectively corresponds to grouping together all mapping profiles connected through at least one shared read mapping. This results in excessive merging of more distinct reference groups and loss of strain-level resolution.

The single-species setting showed similar trends, although the impact of increasing $\epsilon$ was more pronounced because ambiguity is concentrated among highly similar strains within a single species. Overall, the selected default value ($\epsilon = 0.8$) provided a balance between false-positive strain identification reduction and preservation of strain-level resolution across different evaluation settings.

#### Alignment score threshold (high_score_threshold)

The alignment score threshold primarily influenced filtering of weakly supported mappings. Increasing this threshold slightly improved strain identification precision and reduced the number of false-positive strain identifications across all evaluated settings. At the same time, stricter filtering produced small reductions in strain identification recall, indicating that some weakly supported true-positive strain identifications were removed together with false detections.

Read-level classification performance remained comparatively stable across the evaluated threshold range despite reductions in the number of false-positive strain identifications. This suggests that many of the removed false-positive detections were supported by relatively small numbers of reads and therefore had limited influence on overall read-level classification metrics. The observed behavior was consistent across the high-complexity, large-reference-database, and single-species settings.

#### Reference-support thresholds (min_high_score_reads, mid_high_score_reads, min_high_score_fraction)

Thresholds controlling reference reliability produced consistent trends across the evaluated datasets. Increasing the minimum number of high-scoring reads required for a reference to be considered reliable produced the strongest effect, progressively reducing the number of false-positive strain identifications and improving strain identification precision. Similar but weaker behavior was observed for the intermediate-support threshold and the minimum high-score fraction threshold, which together control filtering of moderately supported references during reassignment. Stricter filtering progressively reduced the number of weakly supported candidate references retained during refinement.

These effects were strongest in the high-complexity and large-reference-database settings, where large numbers of ambiguous mappings create substantial opportunities for weakly supported false-positive strain identifications. However, the single-species setting showed a distinct behavior for `min_high_score_reads`, where increasing the threshold substantially improved strain identification precision while preserving strain identification recall. This suggests that stricter support requirements may be particularly beneficial in local strain-resolution scenarios dominated by highly similar genomes, where weakly supported neighboring strains can otherwise accumulate false assignments.

Although stricter support thresholds improved strain identification precision, they also produced moderate reductions in strain identification recall, indicating that overly aggressive filtering may remove low-abundance true-positive strain identifications together with weakly supported false positives. Read-level classification performance remained comparatively stable across most evaluated ranges.

A relatively permissive default value for `min_high_score_reads` was selected to avoid excessive filtering of strains with limited read support. However, the sensitivity analyses indicate that stricter thresholds may be preferable in applications prioritizing maximal false-positive strain identification reduction over preservation of low-abundance detections.

#### Species-level reassignment thresholds (species_min_read_count, species_min_mean_score, species_low_count_cap)

Species-level reassignment thresholds are intended to identify weakly supported species assignments prior to strain-level refinement. Such assignments can arise from ambiguous mappings, low-confidence cross-species alignments, or situations where the true organism is absent from the reference database and reads are distributed across multiple related species. The evaluated parameters therefore define minimum support requirements based on the number of assigned reads and the average alignment score associated with a species.

Species-level reassignment thresholds generally produced comparatively modest performance changes across the evaluated datasets. Increasing the minimum species read count and species mean-score thresholds slightly reduced the number of false-positive strain identifications and modestly improved strain identification precision, while producing only limited effects on strain identification recall and read-level classification performance.

Similarly, varying the low-count cap parameter had relatively minor influence on overall classification behavior, suggesting that the species-level reassignment procedure is comparatively robust to moderate threshold variation once highly unreliable species assignments are removed.

The limited sensitivity of these parameters compared to clustering and reference reliability thresholds likely reflects the role of species-level reassignment as an initial coarse filtering step prior to strain-level refinement. Most strain-level ambiguity therefore remains governed primarily by the clustering and reassignment behavior operating within species-level groups.

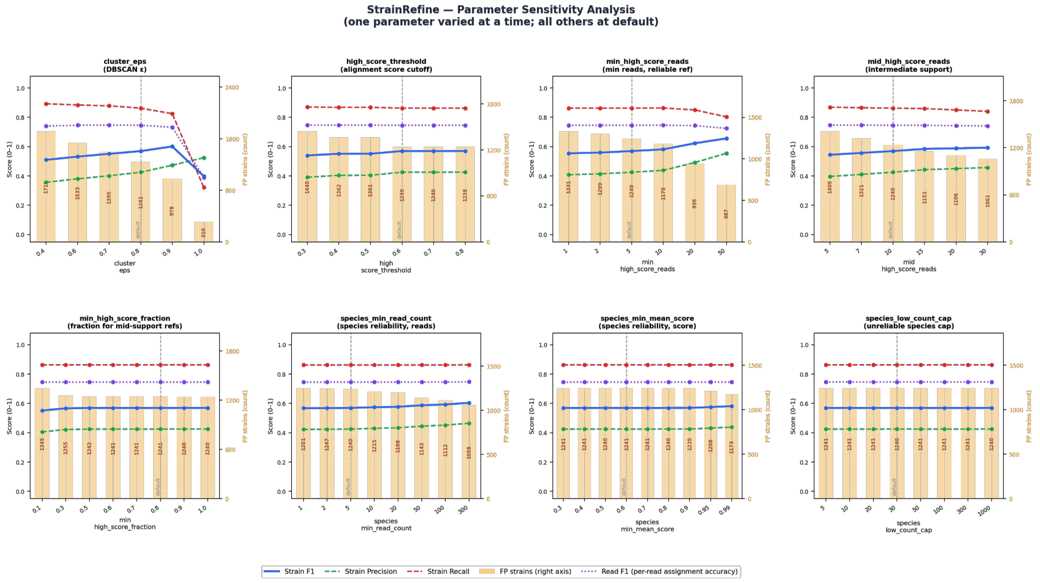


Figure S1: Parameter sensitivity analysis on the high-complexity dataset after MADRe database reduction.

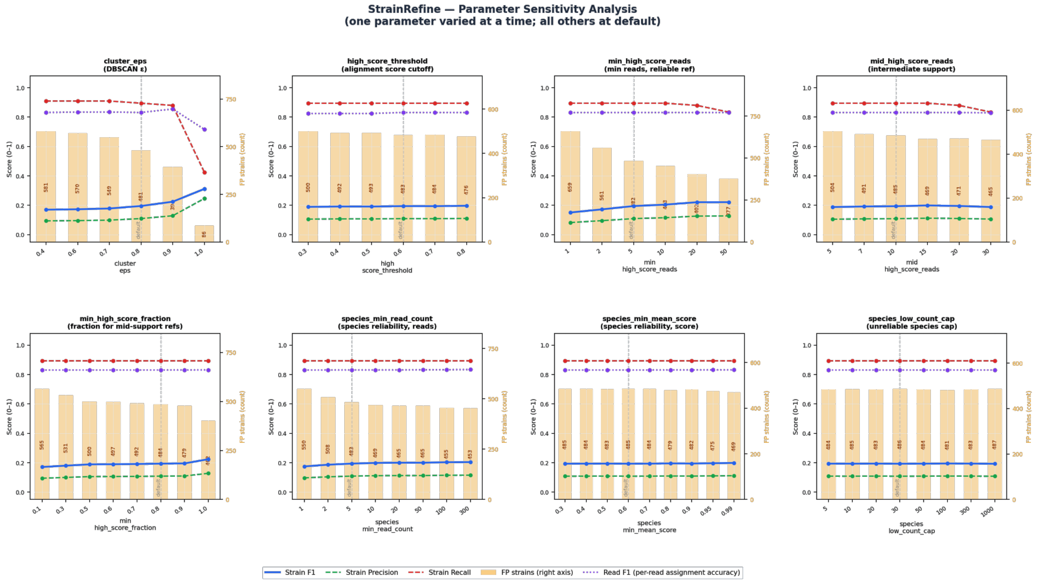


Figure S2: Parameter sensitivity analysis on the large-reference-database setting.

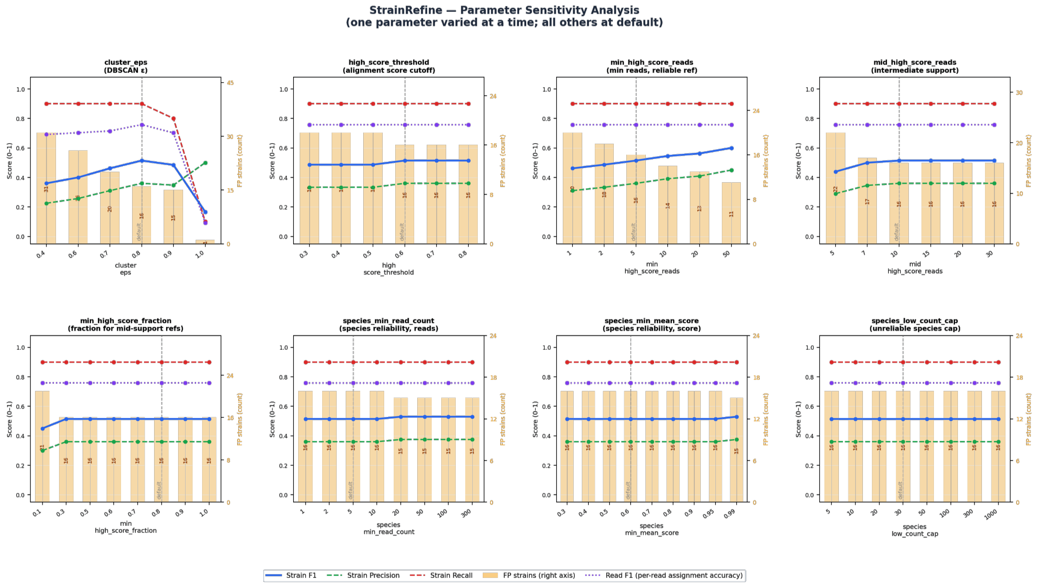


Figure S3: Parameter sensitivity analysis on the single-species dataset containing ten closely related strains.
